# Benchmarking Variational AutoEncoders on cancer transcriptomics data

**DOI:** 10.1101/2023.02.09.527832

**Authors:** Mostafa Eltager, Tamim Abdelaal, Mohammed Charrout, Ahmed Mahfouz, Marcel J.T. Reinders, Stavros Makrodimitris

## Abstract

Deep generative models, such as variational autoencoders (VAE), have gained increasing attention in computational biology due to their ability to capture complex data manifolds which subsequently can be used to achieve better performance in downstream tasks, such as cancer type prediction or subtyping of cancer. However, these models are difficult to train due to the large number of hyperparameters that need to be tuned. To get a better understanding of the importance of the different hyperparameters, we examined six different VAE models when trained on TCGA transcriptomics data and evaluated on the downstream task of cluster agreement with cancer subtypes. We studied the effect of the latent space dimensionality, learning rate, optimizer and initialization on the quality of subsequent clustering of the TCGA samples. We found *β*-TCVAE and DIP-VAE to have a good performance, on average, despite being more sensitive to hyperparameters selection. Based on these experiments, we derived recommendations for selecting the different hyperparameters settings. In addition, we examined whether the learned latent spaces capture biologically relevant information. Hereto, we correlated the different representations with various data characteristics such as age, days to metastasis, immune infiltration, and mutation signatures. We found that for all models the latent factors, in general, do not uniquely correlate with one of the data characteristics even for models specifically designed for disentanglement.

## Introduction

Advancements in sequencing technologies have enabled profiling different “-omics” that revolutionised the understanding of biology. These omics are usually of high dimensionality, which complicates the data analysis. This has sparked a large research interest in dimensionality reduction methods which represent data in a lower-dimensional space while reducing noise and preserving the signal in the data. There are different dimensionality reduction methods that can be categorized into linear and non-linear methods [1] [2]. Selecting an appropriate dimensionality reduction method for an application depends on the structure of the high-dimensional space and the structure of the low-dimensional manifold that we assume that the data belongs to.

Variational AutoEncoders (VAE) are among the most used methods nowadays to embed omics data into a lower dimensional representation. A variational autoencoder is similar to an autoencoder (AE) as they both learn a set of latent variables *z* to encode an input sample *x* and by forcing *z* to be able to reconstruct *x* (i.e.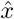). Both VAE and AE are based on an encoder-decoder structure of artificial neural networks (Fig 1A, B). An AE is a deterministic model that is trained by minimizing the reconstruction error of the input data.

**Fig 1.**
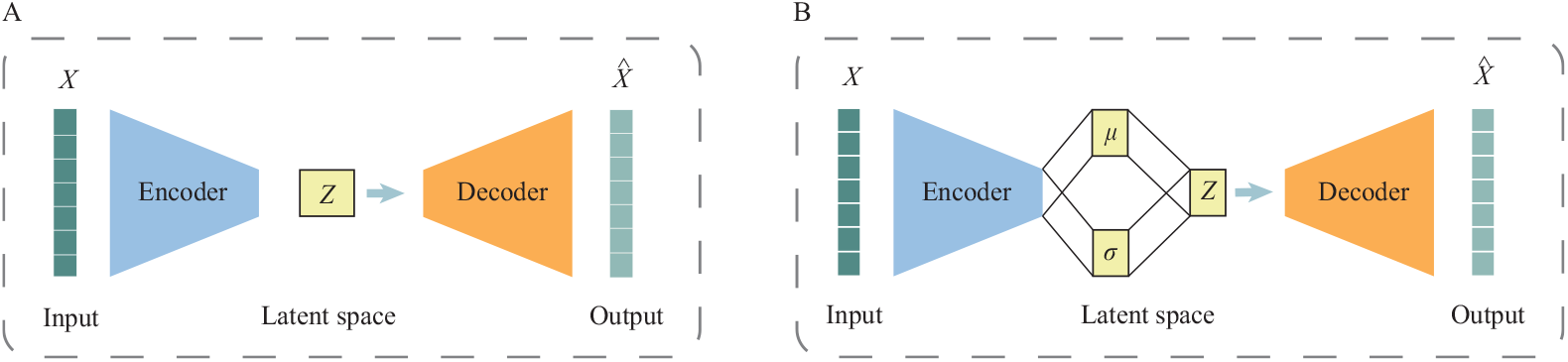
Schematics for autoencoder and variational autoencoder. Both models are based on the encoder-decoder neural network structure to a learn latent space. A) An autoencoder is a deterministic model where *z* is a mapping of the input data. B) A variational autoencoder is a probabilistic model where the mapping *z* is generated by a probability distribution conditioned on the input data.

The VAE differs in that, it learns a probabilistic mapping from *x* to *z* (i.e. a probability distribution *p*(*z*|*x*)) which enables the generation of new data points by drawing samples from this distribution. Calculating this probability distribution *p*(*z*|*x*) is intractable, especially in high dimensional data. To overcome this, Kingma and Welling applied variational inference and neural networks to estimate it by a tractable approximation *q*(*z*|*x*) (see equation 1) [3].

Various VAE variants have been proposed to address different aspects of the VAE formulation and to improve the training of VAEs on specific tasks [4]. One task gaining attention is the interpretability of the learned latent space. Several models have attempted to generate an interpretable latent space by forcing individual latent factors to correspond to specific factors of variation within the dataset, such as biological processes or metadata. Such representations are called disentangled representations [5] [6]. This definition can be generalized to a “set of latent factors” that together encode one independent factor of variation [7]. Different VAE variants have been designed to tackle the disentanglement problem and claim achieving a better performance in learning a more disentangled latent space [8] [9] [10]. Some studies show that the more interpretable the latent space, the better the model is at representing the data [5] [6] [11]. For instance, Way and Greene showed that a VAE can learn a meaningful latent space trained on RNA-Seq data from The Cancer Genome Atlas (TCGA) [12] [13]. Also, VAEs are proven to be useful in several applications, such as predicting drug response [14] and perturbation effects [15]. Using a semi-supervised approach and a VAE, Wei and Ramsey were able to predict response to chemotherapy for some cancer types [16].

Despite these promising results, it is known that VAEs suffer from sensitivity to the hyperparameters, such as the learning rate, the number of hidden layers, the optimizer, the number of neurons in each layer [17] [18] [19]. Although VAEs are getting more and more widely-used, there is a lack of guidelines for selecting training hyperparameters. In addition, there has been no consistent comparison of different VAE models on their ability to learn a disentangled latent space when applied to embed RNA sequencing data of cancer patients [20].

In this paper we study the capability of different VAE models to learn the latent representation of the data and the ability to disentangle this representation. We benchmark the performance of six different VAE variants and four different hyperparameters: dimensionality of the latent space, learning rate, optimizer, and initialization, leading to a total of 3,240 different VAE configurations. The performance was evaluated on the clustering quality of transcriptomic samples in the TCGA dataset which comprises of patients with different cancer types. Moreover, for well-performing hyperparameter configurations, we tested the disentanglement of the learned latent space. Finally, based on our benchmarks, we provide recommendations on selecting VAE models and their hyperparameters when dealing with transcriptomic data.

## Materials and methods

### VAE models

In this paper we selected six different VAE models to benchmark. We focused on VAEs that are designed for learning a disentangled latent space or learning a discrete representation for the input data that is suited for downstream analysis tasks.

#### Vanilla VAE

The VAE model aims to find a probabilistic distribution *p*(*z*|*x*) which maps the input *x* to a set of latent variables *z*. Because *p*(*z*|*x*) is intractable in most cases, we follow Kingma and Willing [3] and approximate it by a distribution *q*(*z*|*x*) with parameters *ϕ* which is approximated by a neural network (encoder). A decoder neural network then tries to reconstruct the input data from the latent variables by learning the distribution *p*(*x*|*z*) with parameters *θ* [3]. VAE achieves this by maximizing the the evidence lower bound (ELBO) which is a lower bound of the data log-likelihood (*p*(*x*)) [21]. This leads to the following loss function which is the negative of the ELBO [3]:

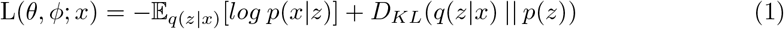

Then, a stochastic gradient variational Bayes estimator [3] is used to minimize this loss with respect to the parameters. The first term in equation 1 corresponds to the reconstruction error which directs the decoder to learn how to accurately reconstruct the input data. The second term is the Kullback-Leibler (KL) divergence between the learned embedding distribution of an input sample and the prior distribution *p*(*z*) which acts as a regularizer for the encoder.

#### *β*-VAE

Higgins et al. introduced the idea of learning a disentangled representation using an adaptation of a VAE called *β*-VAE. This work showed that adjusting the balance between the reconstruction loss and the KL divergence terms can push the encoder to learn a disentangled representation [8]. Thus, this model multiplies the KL term with a hyperparameter *β*, such that the loss function becomes:

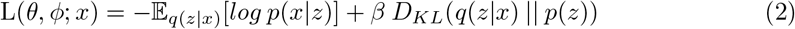

The loss function is similar to equation 1 except for the *β* parameter. When setting *β* > 1, the model is “encouraged” to learn a disentangled representation of the training data [8].

#### *β*-TCVAE

The *β*-Total Correlation VAE, or *β*-TCVAE for short, is an extension of the *β*-VAE model. R. Chen et al. showed that decomposing the loss function of *β*-VAE (i.e. equation 2), and penalizing the total correlation between the latent variables, forces the model to find more statistically independent latent variables [9]. Hence, the loss function becomes:

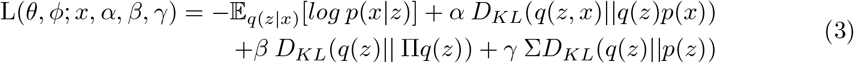

Here, the KL term in equations 1 and 2 is decomposed into 3 different terms. The first term, preceded by *α*, is modeling the mutual information between the data variable and latent variables. The second term, preceded by *β*, is modeling the dependence between the different latent variables and is called the total correlation (TC) term. The last term that is preceded by *γ* is used to prevent each individual latent variable from diverging away from its prior. This work showed that penalizing the total correlation term (i.e. setting *β* in equation 3 to a large positive value) helps the VAE to learn disentangled representations [9]. However, the effect of weighting the three different terms in finding a disentangled latent space is hard to assess.

#### DIP-VAE

Disentangled Inferred Prior VAE (DIP-VAE) learns a disentangled representation by matching the covariance of the prior distribution and the latent distribution. Authors argued that achieving a disentangled representation requires a disentangled prior [10]. The model uses the following loss function:

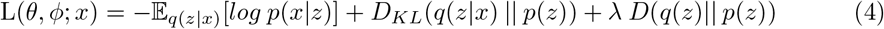

In the last term, *D*(.), denotes a distance metric between *p*(*z*) and the (intractable) *q*(*z*). The authors modeled this distance as the squared difference between the two corresponding covariance matrices. That means that DIP-VAE minimizes the covariance between latent factors, while *β*-TC-VAE minimizes the correlation. This squared difference can in practice be computed in two ways, leading to two sub-variants termed DIP-VAE I and DIP-VAE II (see [10] for more details). Here, we used DIP-VAE II where the latter term is computed as shown in equation 5:

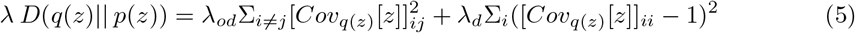

The *λ*_*d*_ and *λ*_*od*_ variables are used to weigh the contribution of the disentanglement objective.

#### IWAE

Importance Weighted AutoEncoder (IWAE) provided a tighter ELBO of the data log-likelihood compared to the vanilla VAE [3]. This was achieved by drawing *K* samples instead of one from the encoder network in order to perform the Monte Carlo estimate of the expectation term of equation 1 [22]. Thus the loss function becomes:

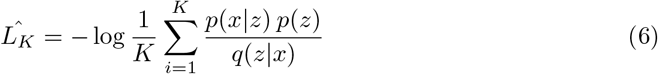

Note that this loss cannot be decomposed to a reconstruction term and a KL divergence term for *K* > 1.

#### Categorical VAE

Categorical VAE (CAT-VAE) made it possible to model a discrete latent space, unlike vanilla VAE that consider continuous Gaussian latent space. CAT-VAE introduced the Gumbel-softmax distribution that is a continuous distribution that approximates the categorical distribution [23]. CAT-VAE can incorporate label information, but for a fair compression between the different models we did not use the labels of the samples in the training of this model. The loss function of CAT-VAE for unlabeled data is shown in equation 7, where we marginalize over all possible labels *y*.

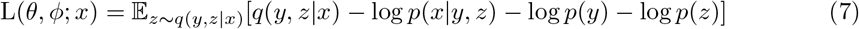

### VAE hyperparameters

We studied the effect of hyperparameters on the training of the aforementioned models. We focused on four types of hyperparameters, whose different settings we explored for every VAE model. We set the *latent dimensions* to be either 10, 20, 30, 50, 100 or 200 factors. For the *learning rate* we used 1e-1, 1e-2, 1e-3, 1e-4, 1e-5 and 1e-6 as step size. For the *initialization of weights* of the encoder and the decoder, we compared the following methods: a standard normal (*N* (0, 1)), Uniform (*U* (0, 1)), Xavier normal, Xavier uniform [24] and Kaming uniform [25]. Finally, we compared three *optimizers* : Adam [26], RMSprop [27] and Stochastic Gradient Descent (SGD) [28].

### Dataset

The models were trained on the TCGA RNA-seq gene expression dataset [13]. The data was downloaded from [29] and samples that have a cancer type label in the meta data were selected [30]. This gave us a total of 11,014 samples from 33 different cancer types. The 5,000 most variable genes across all samples were selected based on the mean absolute deviation (MAD). Each gene was centered and scaled to zero mean and unit variance.

### Evaluation of the effect of hyperparameters

The network architecture was held fixed: The encoder and decoder were made from two fully-connected layers each following the design proposed by Way and Greene [12]. For the encoder, the first layer went from 5000 nodes to 512 nodes, and the second layer went from 512 nodes to the the number of nodes equal to the selected dimension for the latent space. The decoder architecture was the reverse (latent dimension to 512 nodes, and a second layer from 512 nodes to 5000 nodes. For each of the six VAE variants, we ran an exhaustive grid search to evaluate all possible combinations of latent dimensions, learning rate, optimizer and initialization, leading to a total of 540 different setups per VAE variant. To account for variation stemming from the random initialization of the weights, each setup was trained 10 times and we reported the mean and standard deviation of the final loss. Other hyperparameters were left to their default values according to the implementation of [31] and listed in S1 Table.

The dataset was split into a training (70%) and a validation set (30%) stratified per cancer type. We trained the models on the training data for 1000 epochs and applied early stopping if the validation loss did not improve for longer than 5 epochs. The mean validation loss over the 10 random restarts was used as the criterion for evaluating hyperparameter combinations.

We evaluated the ability of the models to cluster the input data in the latent space. To do so, for each hyperparameter combination, the whole dataset was embedded into the latent space (*z*). Then, the embeddings were used to cluster the data using the Leiden community detection algorithm [32]. The neighbourhood graph for the Leiden algorithm was created based on the default settings, using 15 nearest neighbours found according to the Euclidean distance. Then, the Adjusted Rand Index (ARI) was calculated between the found clusters and the known cancer type labels [33].

### Evaluation of disentanglement

We selected the recommended configurations for VAE models (i.e. learning rate, optimizer and initialization) derived from the hyperparameter evaluations (see Results). We then evaluated these configurations on their ability to encode certain data features of interest, solely using one or two latent variables, by calculating the Spearman correlation between a latent variable and a data feature and setting the threshold of correlation at *ρ* = 0.3 which is approximately equivalent to the statistical significance threshold for the correlation using an alpha of 0.05. Data features tested were: age, days to metastasis event, immune infiltration [34], and the presence of either of the mutation signatures SBS 1,2,5,13,15 and 40 determined with exome sequencing [35]. To evaluate the effect of the latent dimension size on disentanglement, we repeated the experiment for latent dimensions sizes in the range 10 to 200 as before.

We measured how many of these 9 data features were encoded by each model, i.e. how many features are correlated with at least one latent factor. Then we measured whether these features are encoded in a disentangled representation, which we defined as being correlated with only one or two latent factors.

All VAEs were retrained using a 70-10-20% split for the training, validation and test set respectively, stratified per cancer type. As these features suffer from missing information across the samples, the Spearman correlation was calculated over all samples (training, validation and test data combined).

### Code availability

The code used in this study can be found on https://github.com/meltager/vae_benchmark.

## Results

### The validation loss does not always reflect downstream performance

We tested the performance of six different VAE models, while varying four different hyperparameters on the TCGA RNA-seq data. To evaluate the learned latent space, we passed all data points through the encoder of each trained model to extract the corresponding embeddings and clustered them using the Leiden algorithm [32]. The models were then evaluated on whether their resulting clustering overlaps with the different cancer types using ARI as evaluation measure. Fig 2A, S1 Fig:S5 Fig, show that despite the correlation between loss and ARI (Spearman *ρ* = − 0.58 for vanilla VAE, S2 Table), models with the same loss can have different ARI values. For example, Fig 2B&C shows the learned embeddings of two hyperparameter combinations that had both a loss of value 1. One of the models generates a clustering resulting in an ARI ~ 0.01 (Fig 2B), while the clustering as a result of other model has an ARI ~ 0.64 (Fig 2C). This implies that the loss value of the VAE does not always reflect the performance of the downstream clustering task.

**Fig 2.**
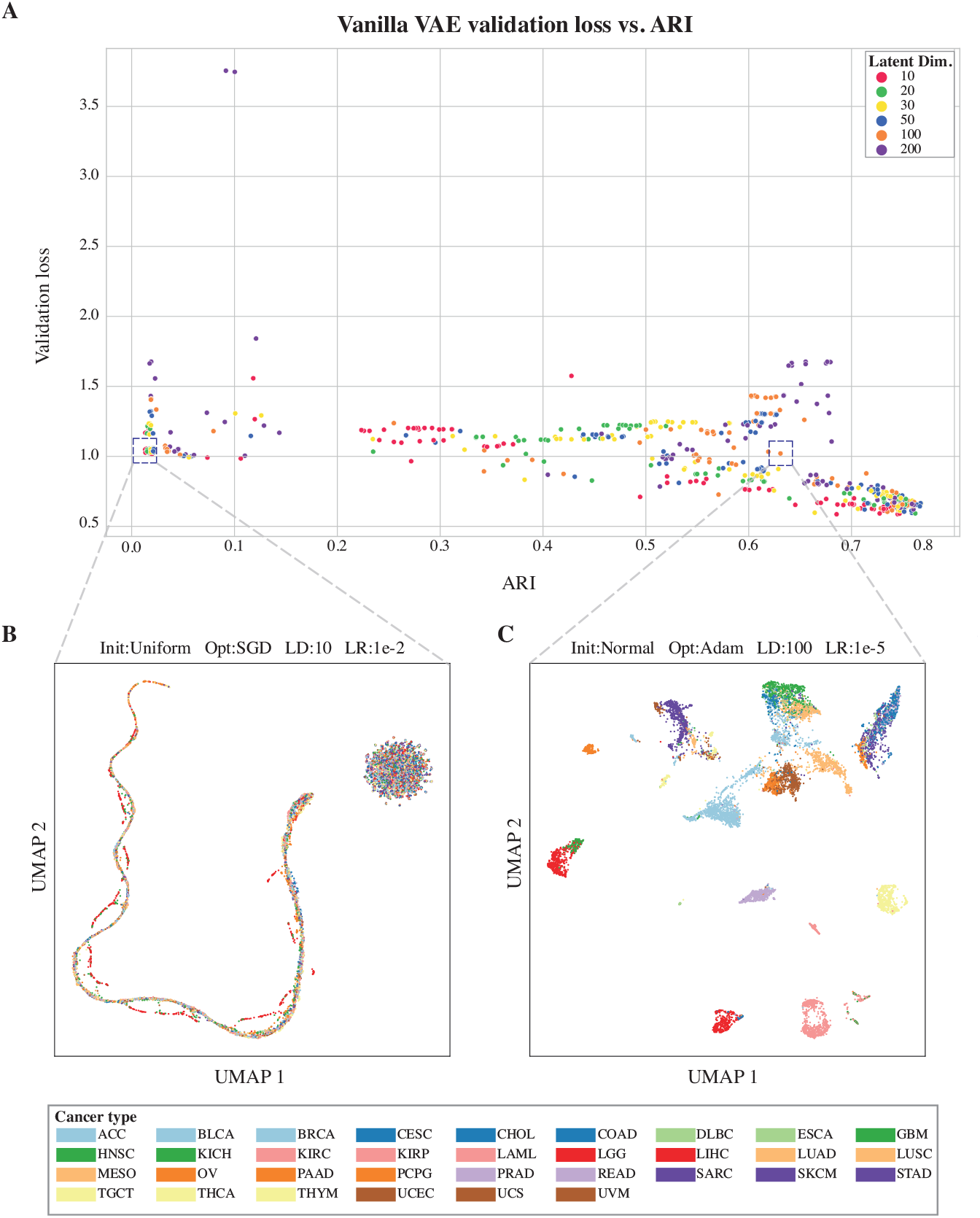
Validation loss does not reflect learned latent space. A) Plotting the validation loss (*y* − axis) vs the ARI (*x* − axis) for the different vanilla VAE hyperparameters configurations. The figure shows a correlation between the validation loss and ARI, however, different configurations with the same validation loss can have different ARI scores. The dots are colored after the latent space dimensions variable. B) UMAP for a configuration for a VAE model that failed to learn how to cluster the data according to cancer types [36]. C) UMAP for a configuration for a VAE model that was able to learn an embedding that can properly cluster the data according to cancer types.

### *β*-TCVAE and DIP-VAE are the best performing models

To assess the performance of the different VAE models, we evaluated the ARI scores across the different hyperparameter configurations. When trained using appropriate hyperparameters, all VAE models could achieve a comparable clustering performance as shown in Fig 3. The boxplots in the figure show the performance of all different hyperparameter configurations used for each VAE model. We found that *β*-TCVAE and DIP-VAE models performed better than the rest on average.

**Fig 3.**
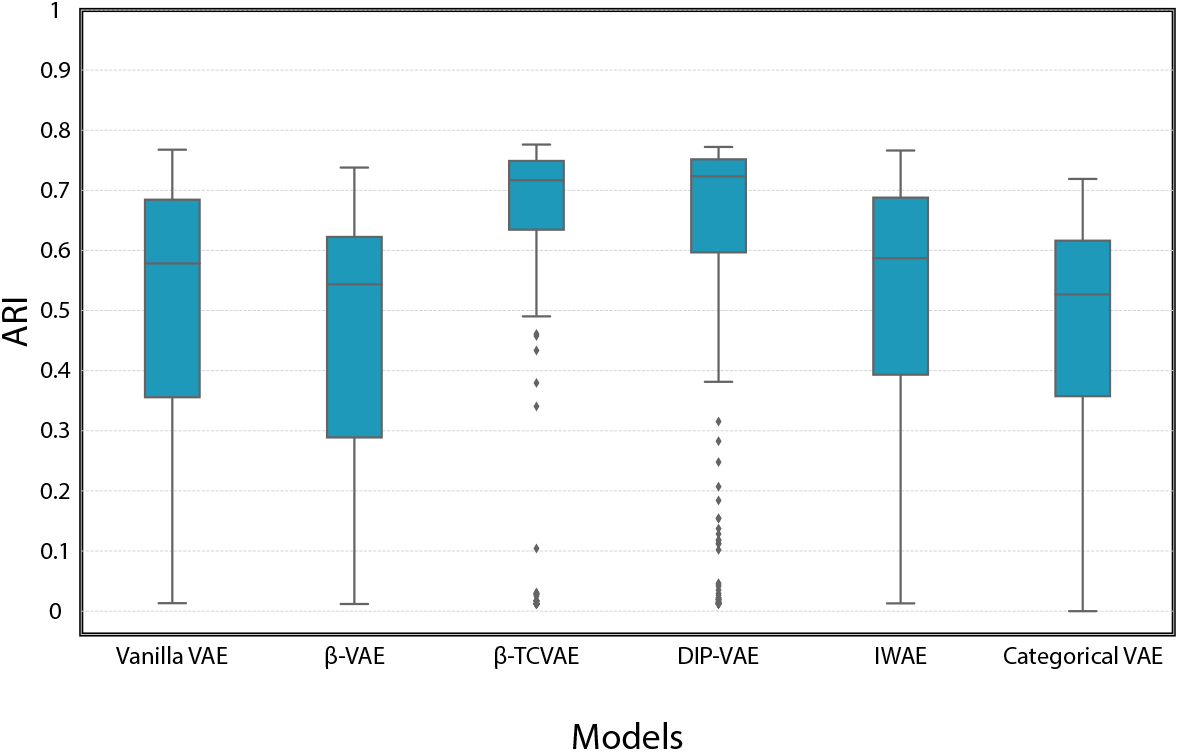
Performance of VAE models in the clustering task. The figure shows the clustering performance (ARI, *y*−axis) compared to the true cancer type using data embedded to the latent space learned by each model (*x*−axis).

### Choice of hyperparameters affects the VAE performance

Next, we tested the effect of each of the four different hyperparameters on the performance of different VAE models in terms of the ARI scores individually. First, we analysed the effect of the number of latent dimensions on the VAE performance (Fig 4A). A small number of latent dimensions resulted in lower performances, mid-range values (i.e. 50 - 100) performed well across all models. When examining the effect of the learning rate on the performance, we found that the smallest and the largest learning rates are not performing as well as mid-range learning rates (i.e. 1e-3, 1e-4) see Fig 4B. Moreover, training *β*-TCVAE with a large learning rate failed because the optimization diverged regardless of the choices for the remaining hyperparameters. In addition, we found that the choice of weight initialization method did not affect performance with the exception of U(0,1) which clearly underperformed the other methods (Fig 4C). For the choice of the optimizer we found that the SGD optimizer on-average results in lower performance, while the Adam optimizer is on average slightly better than RMSprop, see Fig 4D.

**Fig 4.**
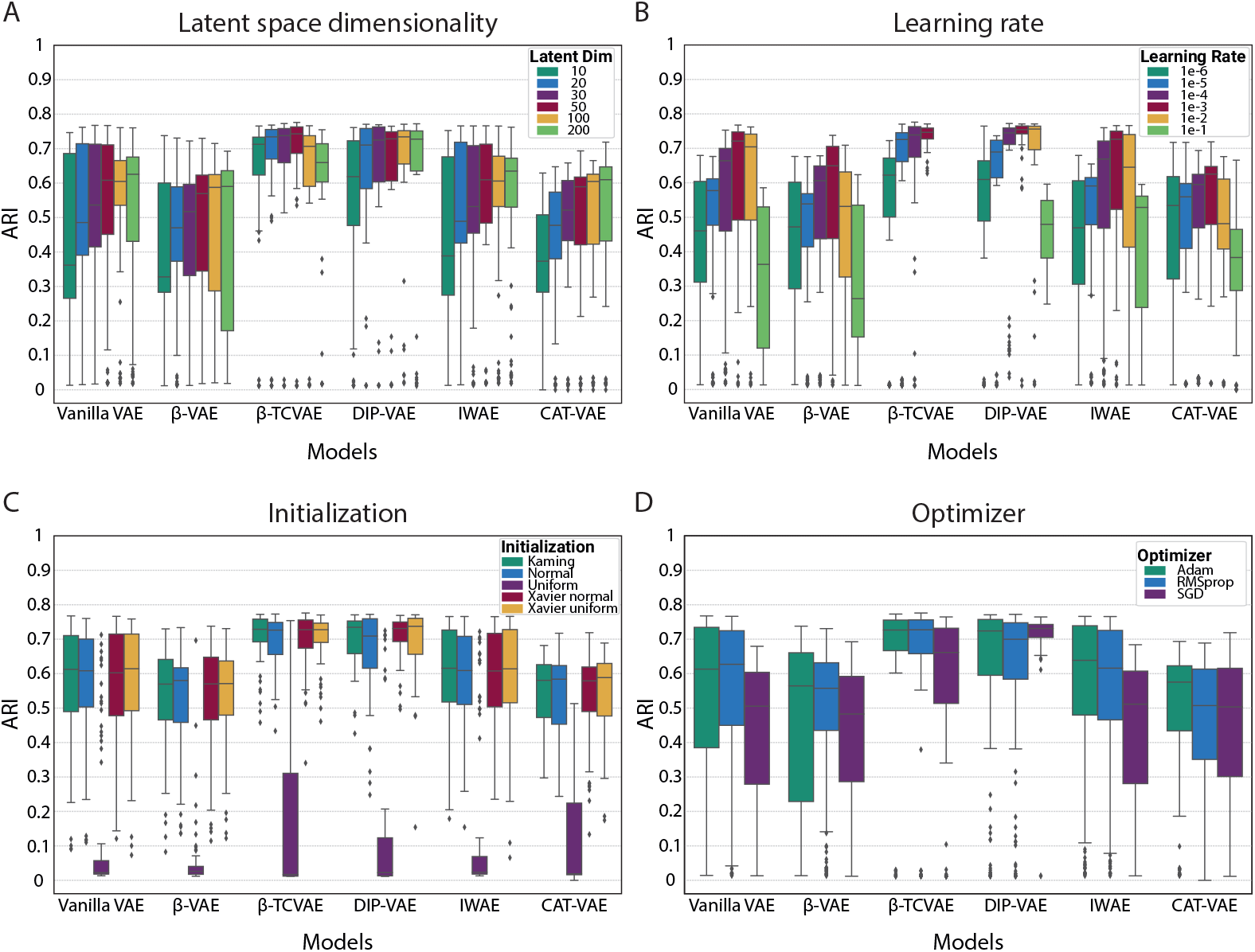
Effect of hyperparameters on different VAE models performance. Each boxplot shows the performance (ARI, *y* − *axis*) of fixing a hyperparameter while varying all others for each VAE model (*x* − axis). The four panels show the four different hyperparameters tested : A) Effect of latent dimensions, B) Effect of learning rate, C) Effect of initialization method, D) Effect of optimizer selection

Finally, we checked the effect of the four different hyperparameters on the viability of the VAE model configuration, i.e. whether the training managed to converge to a solution or (some of) the weight values diverged to infinity. The main cause of failure is the exploding gradient problem, where the network derivatives are getting very large (i.e. explode) leading to an overflow in network update weights, hence failure in updating weights and training of the network. Fig 5 shows all combinations of the different VAE models and hyperparameters and whether they succeeded or not. We found that most failures are coming from the *β*-TCVAE and DIP-VAE models indicating that these two models are more sensitive to the hyperparameter selection. Contrary, Categorical VAE never failed in any hyperparameter combination. The learning rate selection is one of the main causes of failure, the smaller the learning rate the less probable the model to fail. The selection of optimizer and initialization method are less crucial to training failure, while the latent dimensions size is not influencing the model to fail.

**Fig 5.**
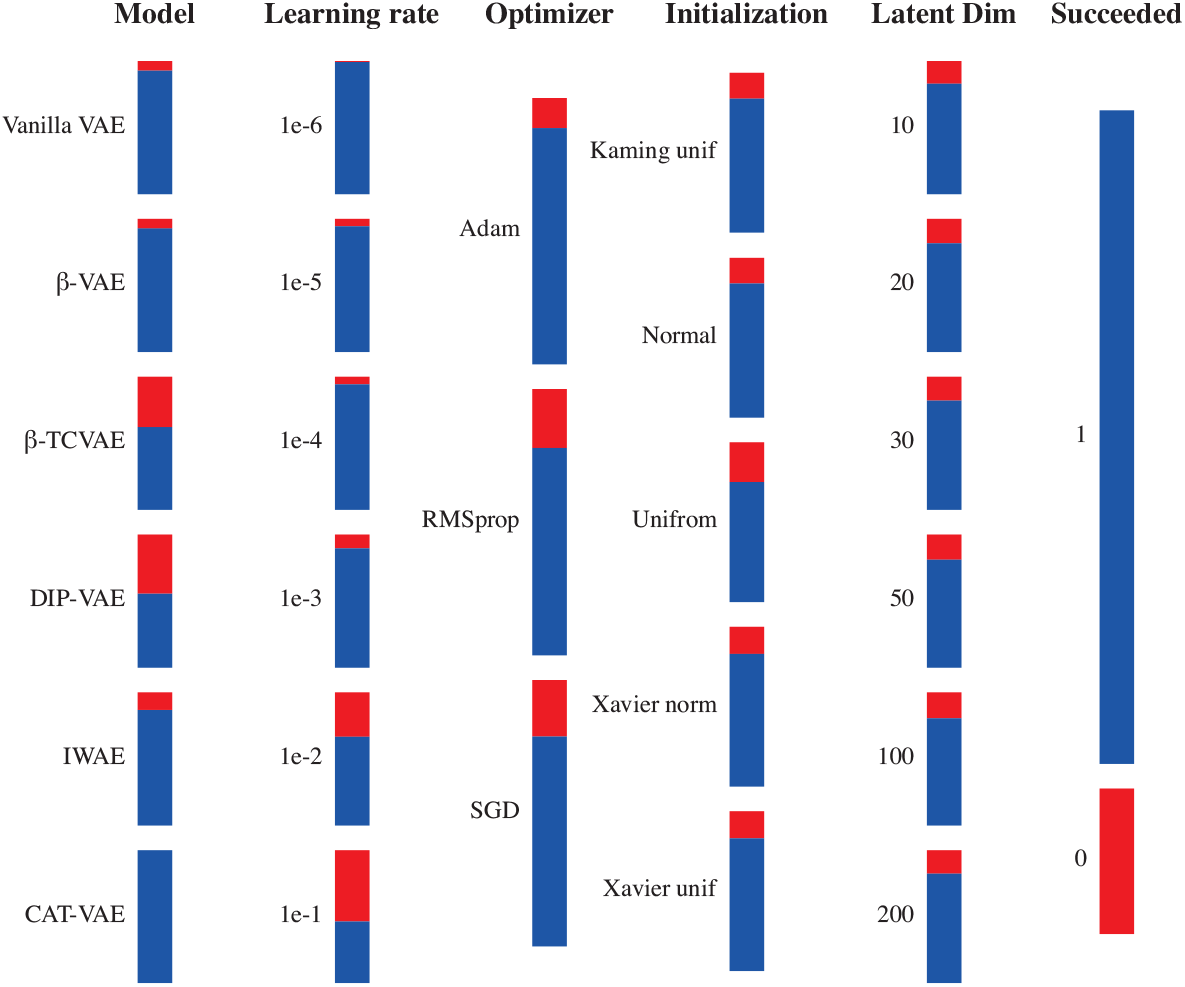
Viability of different VAE models - hyperparameters combinations. Each bar showing all the possible combinations of settings related to mentioned setting. The blue color indicates the number of the succeeded configurations, while the red color indicates the number of the failed ones. Each vertical axis shows all of the 3,240 tested configurations. Each vertical axis, shows the distribution of the failed configurations based on that setting.

### Latent space disentanglement is not trivial to achieve in an unsupervised manner

Based on our benchmark results, we selected the recommended configuration for training VAE models on this dataset. Hereto we used 1e-3 for learning rate, Kaming uniform initialization and Adam as the optimizer. Based on our definition of disentanglement, the results are highly dependent on the size of the latent dimension, thus we tested different latent dimension sizes with the aforementioned configuration.

For example, in Fig 6 we show the Spearman correlation for each latent space factor of the vanilla VAE and *β*-VAE trained with 10 latent dimensions to each of the data features individually. Fig 6A shows that the vanilla VAE resulted in a correlation between data feature immune infiltration and latent space factors 1,4,5,7 and 8. SBS1 is correlated with latent space factors 2,6 and 9. In case of SBS5 only latent space factors 1 and 2 are correlated. SBS13 is correlated with latent space factors 1,2 and 6, and SBS2 is correlated with the same factors in addition to latent space factor 4. These results show that although the VAE has learned to encode at least 5 of the 9 tested features, this encoding is not disentangled, as only SBS5 is associated with only two latent space factors.

**Fig 6.**
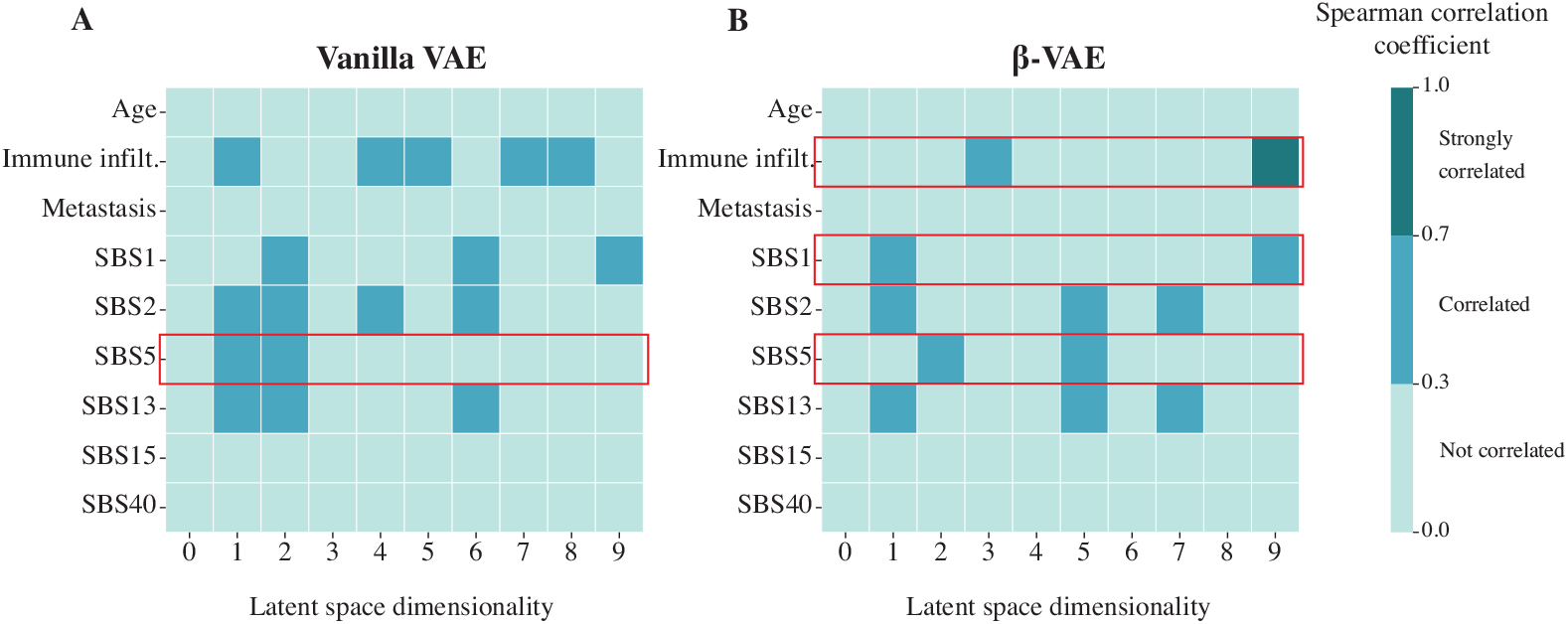
Heatmap showing the Spearman correlation of the latent variables from a 10-dimensional latent space for the vanilla VAE and *β*-VAE with the features. The red highlighting boxes show the disentangled features achieved by a model. A) Vanilla VAE disentangled SBS5 only. B) *β*-VAE disentangled immune infiltration, SBS1 and SBS5

Analogously, the *β*-VAE resulted in a correlation between the same data features but with less number of latent dimensions, Fig 6B. Immune infiltration is correlated with latent space factors 3 and 9. SBS1 is correlated with latent space factors 1 and 9 only. SBS5 is correlated with latent space factors 2 and 5, while SBS2 and SBS13 are correlated with latent space factors 1,5 and 7. Summarizing the *β*-VAE with 10 latent dimensions disentangled 3 data features.

Table 1 shows the disentanglement performances for each VAE model with different latent dimension sizes. The results show that the smaller the latent dimension the better the performance in the disentanglement task. As the number of latent dimensions used for VAE’s increases, more latent space factors start to correlate with the same feature, impacting a model’s disentanglement performance. However, some models can perform better than others for the same latent dimension size. For example, the *β*-VAE and DIP-VAE models that use 10 latent dimensions show the highest number of disentangled latent factors. Only the IWAE model achieved data feature disentanglement when using 200 dimensions for the latent space.

**Table 1.** Number of disentangled data features for different VAE models having different latent dimensions.

| Model \ Latent Dim. | 10 | 20 | 30 | 50 | 100 | 200 |
| --- | --- | --- | --- | --- | --- | --- |
| Vanilla VAE | 1 | 2 | 0 | 1 | 0 | 0 |
| β-VAE | 3 | 1 | 1 | 0 | 0 | 0 |
| β-TCVAE | 1 | 1 | 1 | 1 | 0 | 0 |
| DIP-VAE | 3 | 1 | 1 | 0 | 1 | 0 |
| IWAE | 1 | 1 | 1 | 1 | 1 | 1 |
| CAT-VAE | 1 | 1 | 1 | 1 | 0 | 0 |

To confirm the ability of VAE’s to generalise the disentanglement results, we calculated the correlations using the left out test dataset. S6 Fig shows the correlation between the latent space factors and features as age and immune infiltration features in the test dataset. We can see the similarity in the correlation patterns found in both train and test dataset. From these results we conclude that the disentanglement task is in general difficult for all VAE models and that when selecting the latent dimension size there is a trade-off between disentanglement and downstream performance.

## Discussion

This paper studies settings of different VAE models when applied to cluster cancer patients from their RNAseq profile. We found that the validation loss is not always reflective of the performance on a downstream task that uses the latent space embeddings. Nevertheless, we showed that all VAE variants have the ability to learn a representation of the data that facilitated the downstream task of clustering cancer patients. Despite the fact that *β*-TCVAE and DIP-VAE models had an on-average better performance than others, we can not conclude that they outperform the other models, as all the models could reach a comparable performance based on specific hyperparameter settings. Also, the viability of these two models is too sensitive and susceptible to the hyperparameter selection.

There are multiple possible reasons for the observed inconsistency between the validation loss and the clustering performance. One of them could be the usage of mean square error (MSE) as reconstruction loss, which overemphasizes the effect of outlier samples. RNA-seq technologies suffer from different types of technical noise and artifacts [37], that means some samples could be distorted. These distorted samples do not belong to the actual manifold of the data. Then, the squaring factor in the MSE magnifies these errors, making the VAE tries to adopt to these distorted samples.

Moreover, we evaluate the mean squared error independently for each gene, without taking gene-gene correlations into account. Although this is a standard practice in the literature, explicitly modelling these correlations might lead to a more meaningful evaluation of reconstruction error. Of course, this would significantly increase the complexity and training time of the model, especially for high-dimensional datasets. Also, we think that using a reconstruction loss function that is less susceptible to outliers as Huber loss [38] [39] or quantile loss [40], will help in better approximating the true manifold.

Another possible explanation could be posterior collapse, a common issue with VAE training [41]. Posterior collapse occurs when the posterior distribution of one or more of the latent variables (*q*(*z*|*x*)) becomes equal to its prior (*p*(*z*)). In other words, the encoder output is random and does not depend on the input sample, so that the collapsed latent features do not encode any meaningful information about the input. When this occurs during training, a flexible-enough decoder can still learn to (partially) reconstruct the input by ignoring the collapsed latent features and/or by overfitting to the encoder’s output. This leads to a relatively low reconstruction loss, despite the fact that the latent features are not a meaningful representation of the data.

One of the main contributions of this paper is that we can provide recommendations on hyperparameters settings when dealing with cancer bulk RNA data. Our results show that the selection of hyperparameters greatly influences the performance of the VAE, although this might not be surprising. We recommend using latent space of 50 - 100 dimensions. This fits our intuition that too many factors may cause overfitting on the training data, whereas too few factors would not be able to capture the true manifold. Nevertheless, we found that learning disentangled representations in an unsupervised manner is very hard when using that many latent factors and a smaller latent dimension size is preferred if interpretation is important. For the learning rate, we recommend using learning rates between 1e-3 and 1e-4. Large learning rates push a model over the optimum resulting in an oscillating behavior or even make the training fail, whereas low learning rates slow down the learning process tremendously and can get more easily stuck in local minima. For the initialization methods, the uniform methods are not favorable in the deep learning field as the gradient is the same for many nodes, which makes it hard during the training for weight update [42] [43]. The remaining weight initialization strategies we tested, included Kaming uniform (default in PyTorch) and Xavier normal (default in Keras) initialization distributions. All these settings resulted in a comparable performance. The SGD optimizer is found to underperform in the other settings. The Adam optimizer is becoming a *de facto* standard, and is widely used in the deep learning, as it is faster and requires less memory to run. Our results show that the Adam optimizer outperforms SGD, and does slightly better than RMSprop. These results are in line with the results by Kingma and Ba for image data [26].

Interpretation of machine learning and deep learning models is crucial for their eventual adoption into clinical practice, but this still remains challenging. If (some of) the learned latent factors directly correspond to specific interesting aspects of the data, such as biological processes or important covariates, it would improve the interpretability and therefore the value and potential usage of the VAE models. Our experiments in measuring the disentanglement of the latent factors showed that all VAE models only moderately capture the characteristics tested in the TCGA dataset and that disentanglement often comes at the cost of less good clustering in z-space. This even holds for the models that were specifically designed for learning disentangled representations. In our experiments, VAE models were able to correlate the same latent factors for both SBS2 and SBS13 which are known to occur in the same samples. These mutational signatures are connected to the activity of the AID/APOBEC family of cytidine deaminases and the activity of the APOBEC enzyme [44] [45]. On the other hand, we used the age as a negative control on the disentanglement experiment, as the age is not a source of variation in the data. Thus, all VAE models did not correlate (*ρ* > 0.3) any factor with age, which shows that the VAE models did not randomly find a correlation with this feature. § Again our results align with the theoretical proof of the impossibility of achieving complete disentanglement with completely unsupervised learning [6].

Although the promise of disentanglement with VAEs seems unfulfilled, there are three promising alternatives to force models to learn disentangled representations. The first uses semi-supervised VAE models, where known values of the factors to be disentangled are used to guide the VAE training [46] [47]. The second, stemming from computational neuroscience, imposes biologically-inspired constraints on the weights that enhance selectivity of neurons thereby leading to disentanglement [48] [49]. The third approach relies on the existence of another observed variable, which can be harvested to transform the VAE into non-linear Independent Component Analysis [50]. For example, for the TCGA data this additional variable can be the mutation or methylation profiles of the tumor samples. Further research is needed to validate the utility of these ideas on -omics data.

One limitation of our study is that not all hyperparameters combinations were tested for all models. The effect of hyperparameters weighting the disentanglement terms differ between the different VAE models. We did not study the effect of these hyperparameters on the downstream task as well as the disentanglement task. We decided not to do so because this would result in a unfair (unsystematic) comparison between models. Yet, an important hyperparameter, the number of hidden layers and nodes in a hidden layer, we also did not further explore merely because this would increase the space of models immensely.

In conclusion, we benchmarked several VAE variants on transcriptomics data and studied their learned latent spaces in terms of data clustering and disentanglement. Despite a general difficulty to achieve good disentanglement, we found that *β*-TC-VAE and DIP-VAE tend to perform best in both tasks, although their training can more easily become unstable when using inappropriate hyperparameters.

## Supporting information

**S1 Fig.**
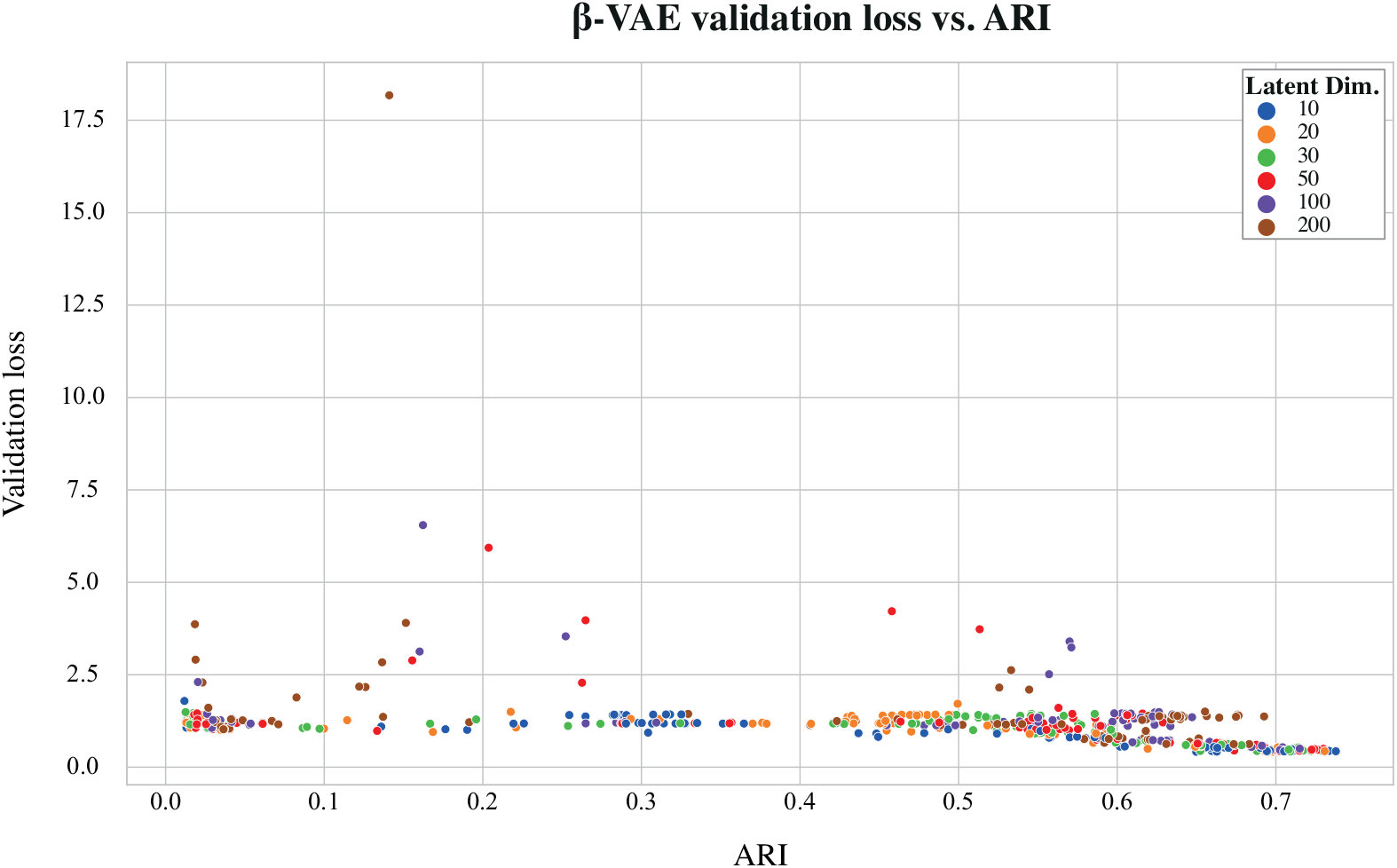
*β*-VAE validation loss vs ARI. Scatter plot for the validation loss of different hyperparameters configurations of *β*-VAE vs ARI. Each dot is a different configuration, and they are colored after the latent space dimensions variable.

**S2 Fig.**
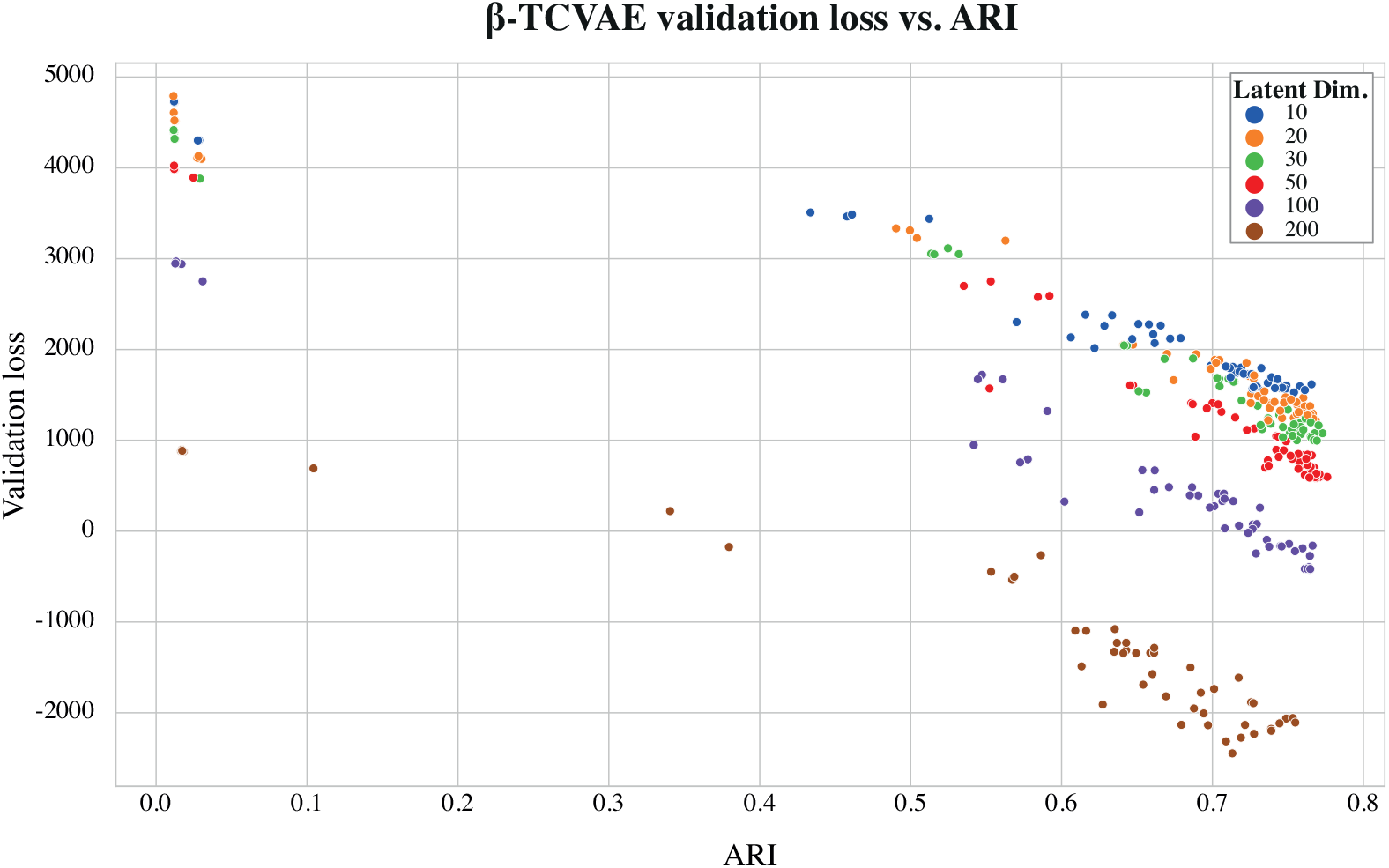
*β*-TCVAE validation loss vs ARI. Scatter plot for the validation loss of different hyperparameters configurations of *β*-TCVAE vs ARI. Each dot is a different configuration, and they are colored after the latent space dimensions variable.

**S3 Fig.**
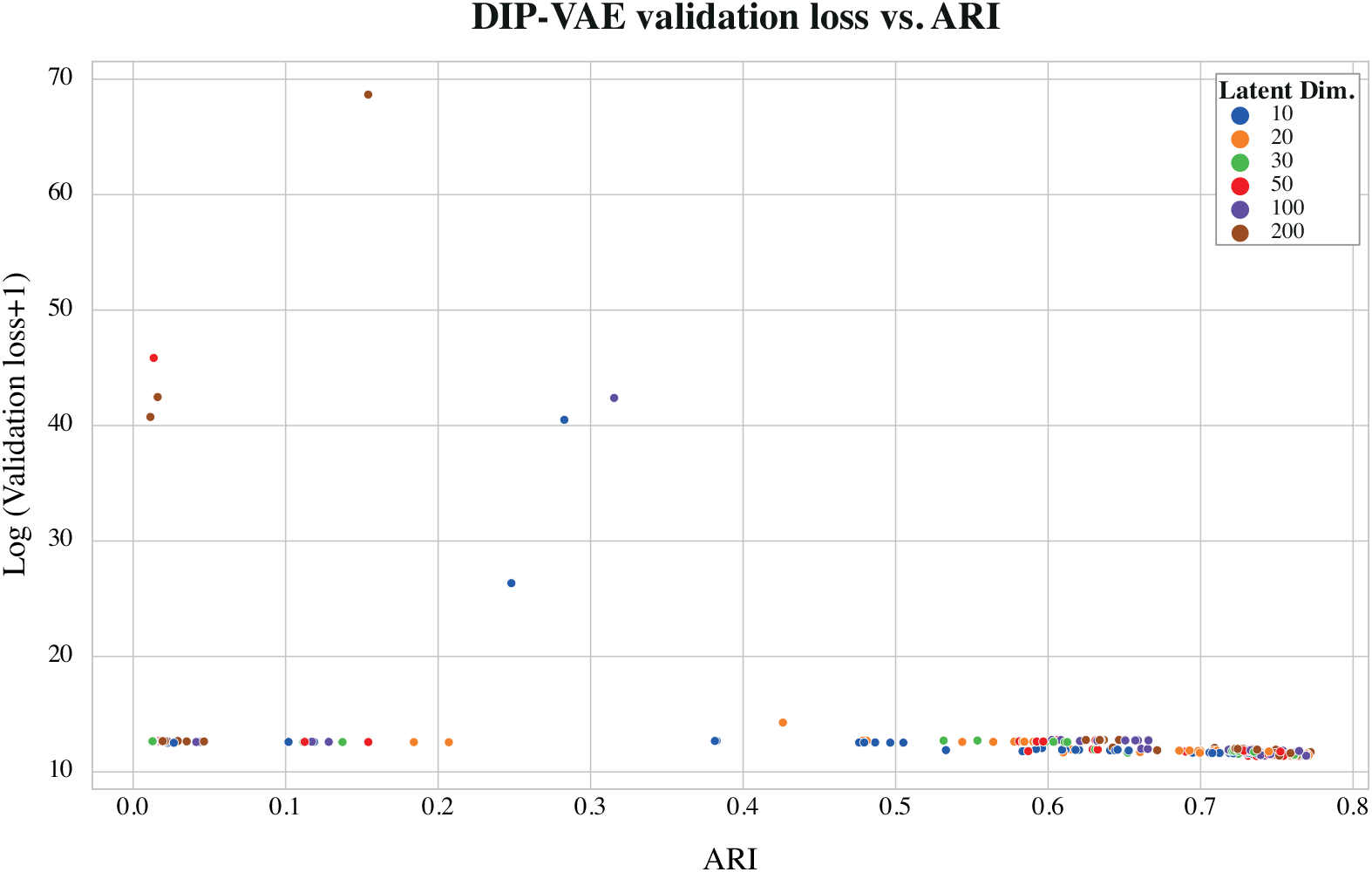
DIP-VAE validation loss vs ARI. Scatter plot for the log of one plus validation loss of different hyperparameters configurations of DIP-VAE vs ARI. Each dot is a different configuration, and they are colored after the latent space dimensions variable.

**S4 Fig.**
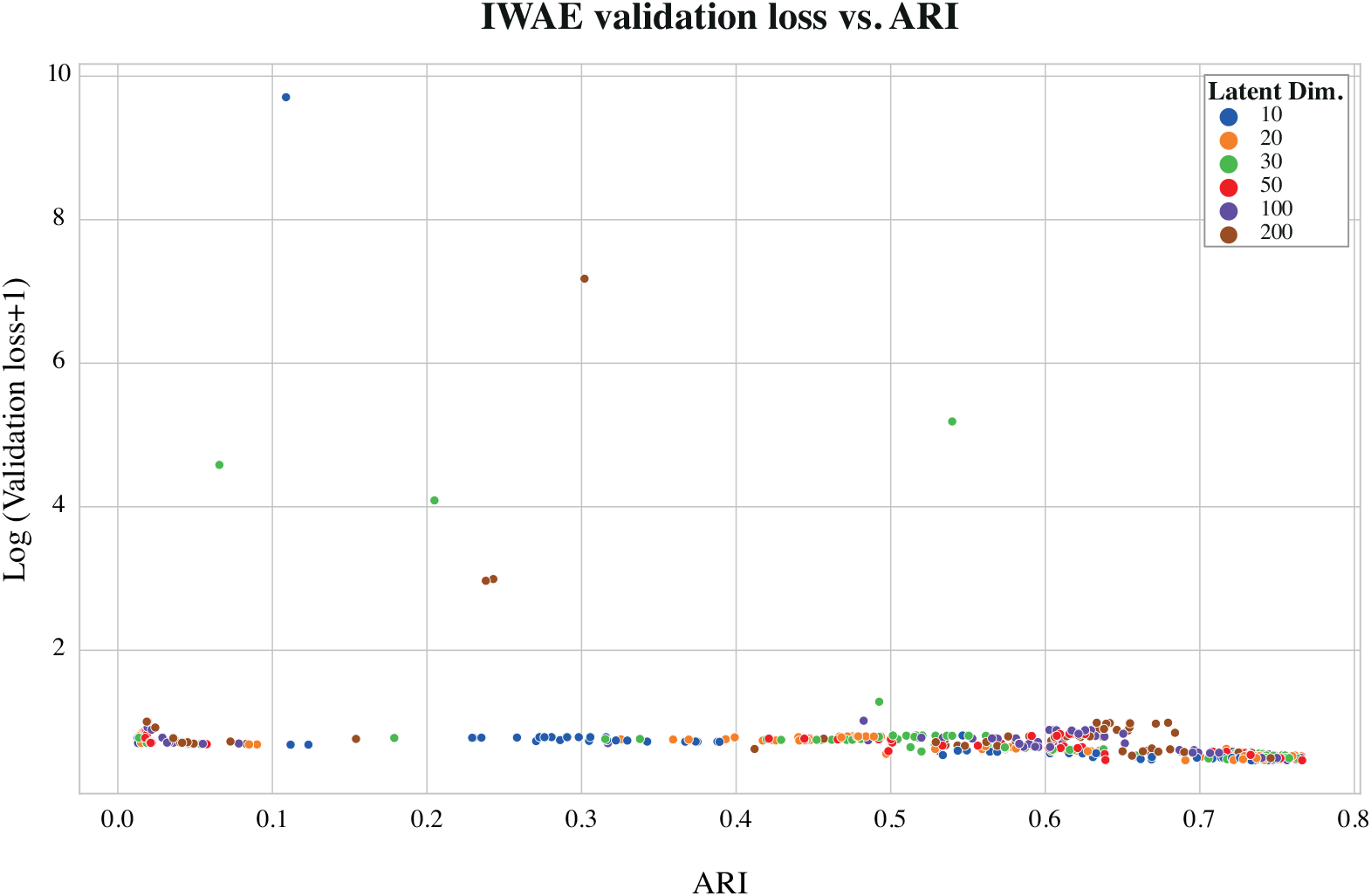
IWAE validation loss vs ARI. Scatter plot for the log of one plus validation loss of different hyperparameters configurations of IWAE vs ARI. Each dot is a different configuration, and they are colored after the latent space dimensions variable.

**S5 Fig.**
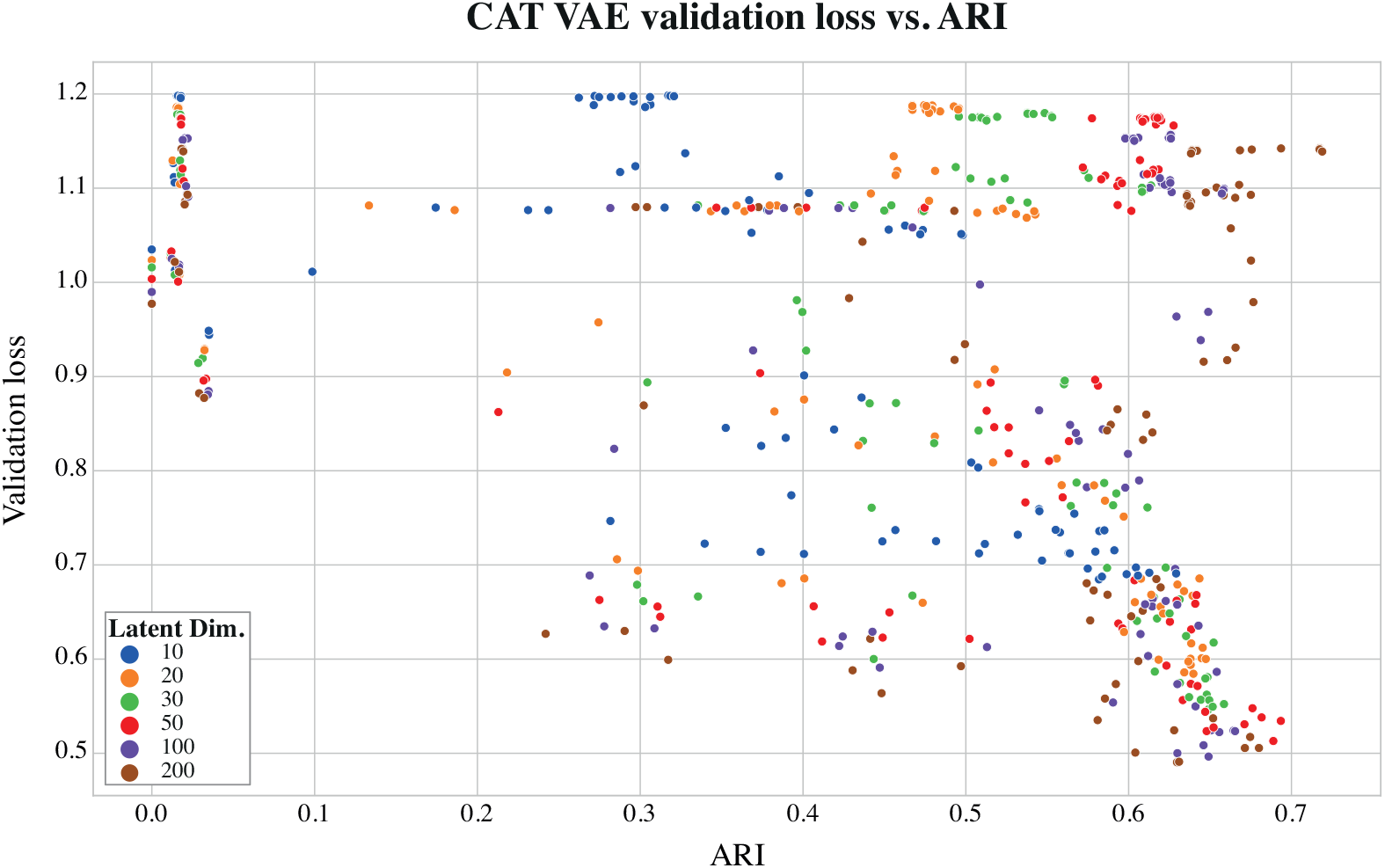
CAT-VAE validation loss vs ARI. Scatter plot for the validation loss of different hyperparameters configurations of CAT-VAE vs ARI. Each dot is a different configuration, and they are colored after the latent space dimensions variable.

**S6 Fig.**
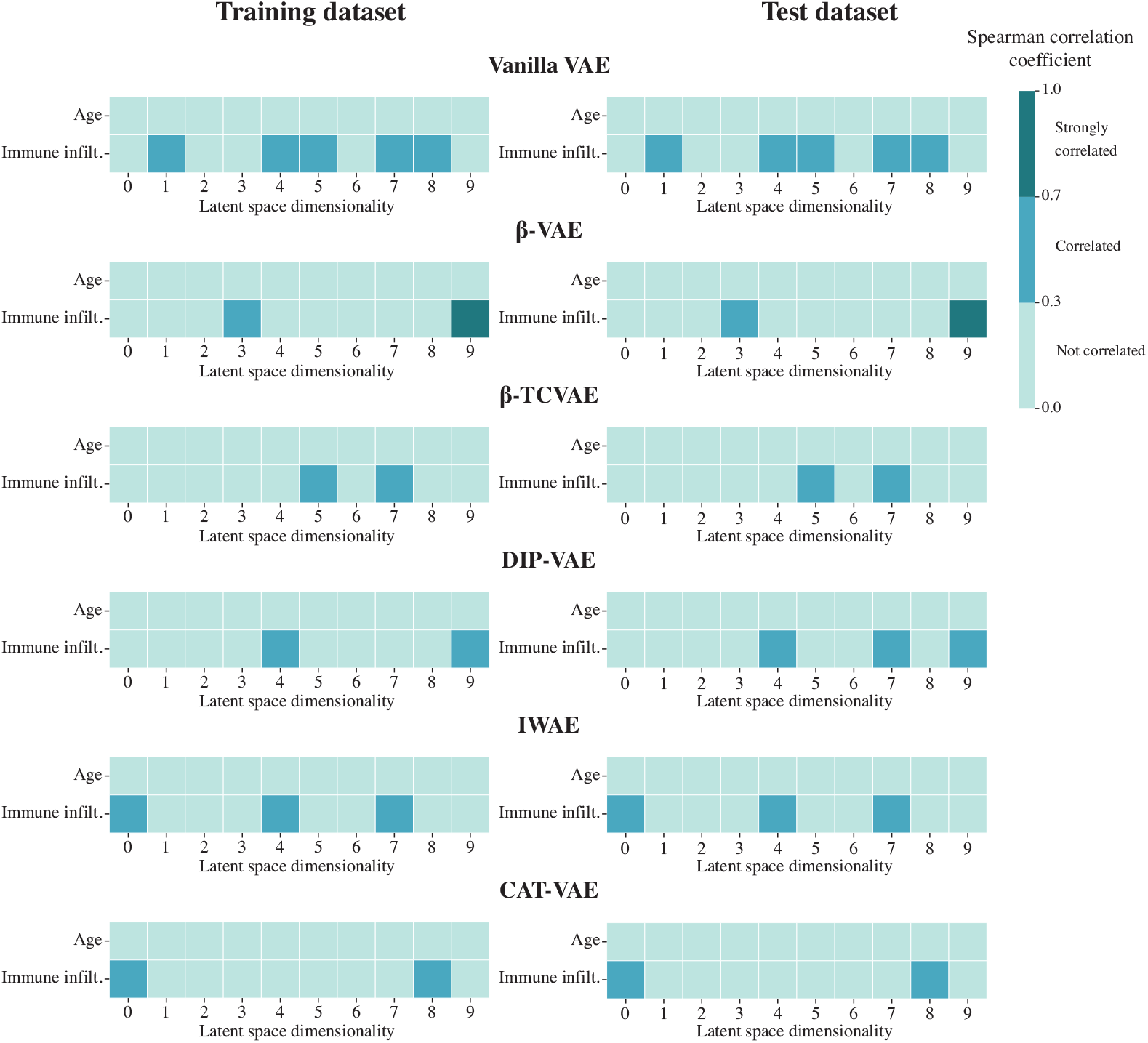
Heatmaps displaying the correlation pattern similarity between latent space dimensionality and data features for both training and test datasets in the VAE models. The figure shows the correlation of age and immune infiltration features with the latent space factors. The left side shows the correlation in the train dataset while on the right side is the test dataset. The figure list all the tested VAE models in the paper.

**S1 Table.** VAE models hyperparameters. A listing of the hyperparameters that were held constant throughout the study. The values were set according to the implementation of [31].

| Model | Parameter | Value |
| --- | --- | --- |
| Common parameters | Batch size | 64 |
|  | Scheduler gamma | 0.95 |
|  | Weight decay | 0.0 |
|  | Maximum epochs | 1000 |
| β-VAE | $\beta$ | 6 |
| β-TCVAE | $\beta$ | 6 |
| | $\alpha$ | 1 |
| | $\gamma$ | 1 |
| DIP-VAE | $\lambda_d$ | 0.05 |
| | $\lambda_{od}$ | 0.1 |
| IWAE | K | 5 |

**S2 Table.** Spearman correlation between different models validation loss and ARI. The absolute rounded Spearman correlation between all the different configurations tested for each model and the ARI values achieved by this model in the downstream task.

| Model | $\rho$ |
| --- | --- |
| Vanilla VAE | 0.58 |
| β-VAE | 0.49 |
| β-TCVAE | 0.43 |
| DIP-VAE | 0.86 |
| IWAE | 0.58 |
| CAT-VAE | 0.3 |

